# Subanesthetic Ketamine Disrupts Predictive Signaling in the Prefrontal Cortex

**DOI:** 10.1101/2025.11.24.690250

**Authors:** Benjamin W. Corrigan, Megan Roussy, Rogelio Luna, Roberto A. Gulli, Jeffrey D. Schall, Adam J. Sachs, Lena Palaniyappan, Julio C. Martinez-Trujillo

## Abstract

Corollary discharge (CD) signals allow the brain to predict and suppress the sensory consequences of its own actions, providing stability to perception and thought. Disruption of these predictive mechanisms has long been hypothesized to contribute to the disorganization of experience in schizophrenia, yet direct circuit-level evidence has been lacking. Here, we show that ketamine, a dissociative N-methyl-D-aspartate receptor (NMDAR) antagonist, at subanesthetic doses, selectively disrupts CD signaling in the lateral prefrontal cortex (LPFC)—a region thought to be the seed of mental representations and one of the most affected areas in schizophrenia. We recorded activity from 1,342 neurons in LPFC areas 8a and 9/46 of macaques performing a visuospatial working-memory task in a virtual environment, before and after subanesthetic ketamine administration. Ketamine impaired performance and increased overall firing rates but markedly suppressed saccade-related responses carrying CD signals. This led to decreased discriminability of eye movement signals. This finding links two major ideas in neuroscience research: the role of disrupted glutamate signaling and the failure of the brain’s predictive models. It provides evidence for how these mechanisms may interact in the prefrontal cortex to disturb the sense of reality.

## Introduction

Schizophrenia is a severe psychiatric disorder characterized by disrupted perception, cognition, and behavior. While its pathophysiology remains poorly understood, mounting evidence suggests dysregulation of glutamatergic neurotransmission plays a central role (Krystal et al., 2003; Moghaddam & Javitt, 2011). Ketamine, a dissociative agent used clinically and recreationally, has been classified as an NMDA receptor antagonist and has been linked to schizophrenia. Indeed, at subanesthetic doses, ketamine induces schizophrenia-like symptoms in healthy individuals and exacerbates symptoms in patients, making it a valuable tool for investigating the disease’s pathophysiological mechanisms (Anticevic et al., 2015; Krystal et al., 1994). However, there is currently a lack of studies in primates that examine circuit-level mechanisms in relation to the main theories of schizophrenia.

Among the well-documented deficits in schizophrenia is impaired corollary discharge (CD)—a neural signal that helps distinguish self-generated sensations resulting from our actions from externally generated sensory input (Ford & Mathalon, 2012). CD impairment is particularly evident in the oculomotor system, where patients exhibit deficits in tasks requiring the integration of eye movement information with visual processing (Thakkar et al., 2015). For example, in a double-step saccade task, patients demonstrate a reduced ability to compensate for the first eye movement, likely due to impaired generation of CD signaling when planning the second, reflecting disrupted internal monitoring of eye movements (Richard et al., 2014; Thakkar et al., 2015).

Concurrent with these CD deficits, patients with schizophrenia show altered activity in the lateral prefrontal cortex (LPFC), a region critical for cognitive control, working memory, and saccade-related processing (Goldman-Rakic, 1995; Weinberger et al., 1986). The LPFC has also been shown to encode mental representations of the visual space during virtual reality tasks in space center coordinates (Busch et al., 2024; Roussy et al., 2022). The LPFC receives extensive projections from oculomotor regions and exhibits saccade-related activity, suggesting that extraretinal signals carrying the parameters of eye movements are integrated within the LPFC circuitry (Bullock et al., 2017; Funahashi et al., 1991; Selemon & Goldman-Rakic, 1988). Such signals may inform the LPFC in advance about changes in the retinal images produced by saccades. A potential source of saccade-related neural signals in the LPFC is CD, which facilitates the integration of motor commands with sensory feedback (Subramanian et al., 2019). It is reasonable to hypothesize that disrupted LPFC processing due to CD abnormalities would affect the ability to predict the consequences of saccades, producing a mismatch between our internal models of reality and current sensory inputs. If sustained, the latter may induce dissociative states and schizophrenia symptoms (Rösler et al., 2015; Thakkar & Rolfs, 2019).

Recent studies suggest that subanesthetic ketamine replicates key aspects of schizophrenia, including thalamic dysconnectivity and working memory impairments (Abram et al., 2022; Roussy et al., 2021). However, the specific effects of ketamine on post-saccadic neural responses linked to CDs in the LPFC microcircuitry remain unexplored. Investigating these effects could provide critical insights into the neural basis of perceptual instability in schizophrenia, as well as validate the administration of ketamine as a translational model for studying these deficits.

Here, we recorded neural activity from LPFC areas 8a and 9/46 in macaques performing a working memory task in a virtual environment before and after ketamine administration. The animals’ gaze was unconstrained, allowing them to inspect the visual scene freely. We examined both behavioral and neural correlates of saccadic eye movements to determine whether ketamine influences saccade-related activity potentially linked to CD signals. We found that ketamine caused an overall increase in neuronal activity. However, in neurons encoding saccade direction, ketamine specifically decreased their activity. Such effects were also present at the level of neuronal populations and began to recover 30-60 minutes after ketamine administration. We used state space analyses and demonstrated that ketamine causes a shift in a low-dimensional neural representation of the saccade command, which affects the discriminability of saccade parameters. Such a shift also begins to recover 30-60 minutes after administration.

## Results

We trained two male macaques in a spatial working memory task set in a virtual environment that they navigated through using a joystick (**Fig. 1a,b**). Each session consisted of a pre-ketamine period of at least 28 trials, followed by a subanesthetic dose of ketamine (0.25-0.4 mg/kg), after which the animal was presented with the task again for 30 minutes. During these sessions, we recorded neural activity in the left lateral prefrontal cortex using two 96-channel Utah microelectrode arrays (**Fig. 1c**). The animals were able to perform the task in the pre-ketamine period (mean performance 74% (SD 20%), and in the post injection period, performance 35% (SD 24%), but with significantly reduced accuracy (**Fig. 1d-f**, see also Roussy et al., 2021). During sessions where saline was injected instead, there was no significant difference between pre-77% (SD 17%) and post-injection performance 75% (SD 18%).

**Figure 1.**
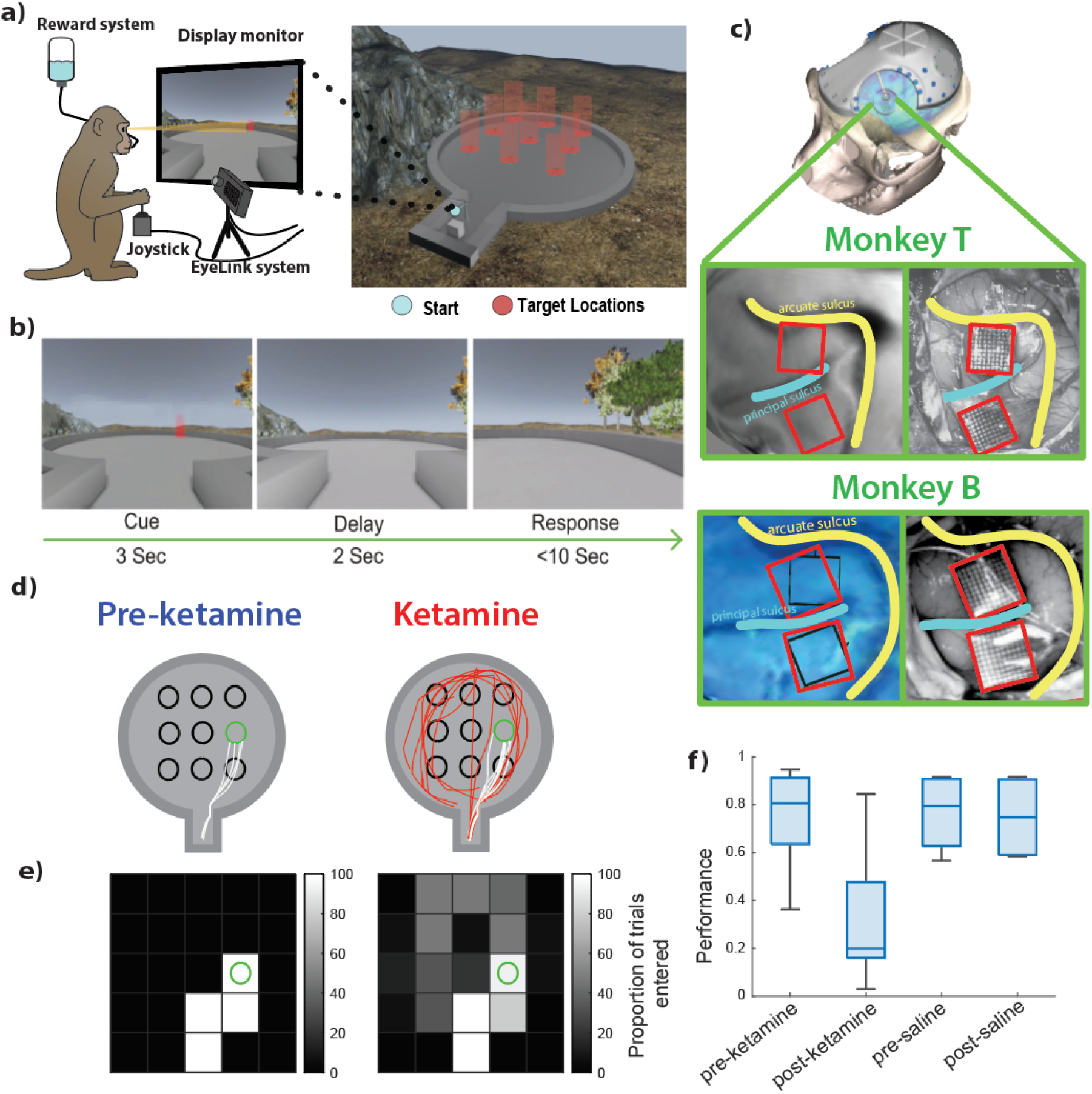
Working memory task performance and recording locations. a) task set-up: monkey seated in front of a screen and used a joystick to navigate the virtual environment (right) while eye position was tracked and juice reward was delivered after correct trials. b) Trial sequence where movement was prevented for the Cue and Delay periods, after which, the monkey navigated freely to the cued location. c) Recording locations of two Utah arrays on either side of the caudal aspect of the left principal sulcus, targeting areas 9/46 and 8a. d) example trajectories of correct (white) and incorrect (red) trials during pre-injection (left) and post-injection (right) to the middle-right target. e) Visitation map where each pixel indicates the proportion of trials where that pixel was entered, with more exploration of the map on post-ketamine trials. f) Box plot of session performance on pre- and post-ketamine trials compared to control saline sessions, where boxes indicate middle quartiles of the distribution, and whiskers indicate full range of performance.

### Effect of ketamine on saccade kinematics

We compared the parameters of saccades recorded during pre- and post-injection periods. We only analyzed saccades with 1) a large horizontal component (tolerance of 30° from the horizontal meridian), 2) with amplitudes of 5-20 degrees of visual angle, and 3) covering a similar area during pre- and post-ketamine administration (**Fig. 2a,b**). The main sequence of saccades describes the relationship between saccade amplitude and peak velocity (Crawford et al., 2003) and resembles an exponential grow function during both pre-injection and post-injection saccades (**Fig. 2c**). We found a small but significantly higher amplitude and velocity for saccades made before compared to after injection (Kolmogorov-Smirnov test. amplitude: *D*(14285) =.03, *p* < 0.05, direction: *D*(14285) =.21, *p* < 0.05, **Fig. 2d,e**). We hypothesize such a decrease may be due to the mild sedative effects of ketamine as reported for other sedatives (Busettini & Frölich, 2014). Indeed, in macaques, ketamine has been shown to decrease the velocity for horizontal saccades of magnitudes > 10° (Pouget et al., 2025).

**Figure 2.**
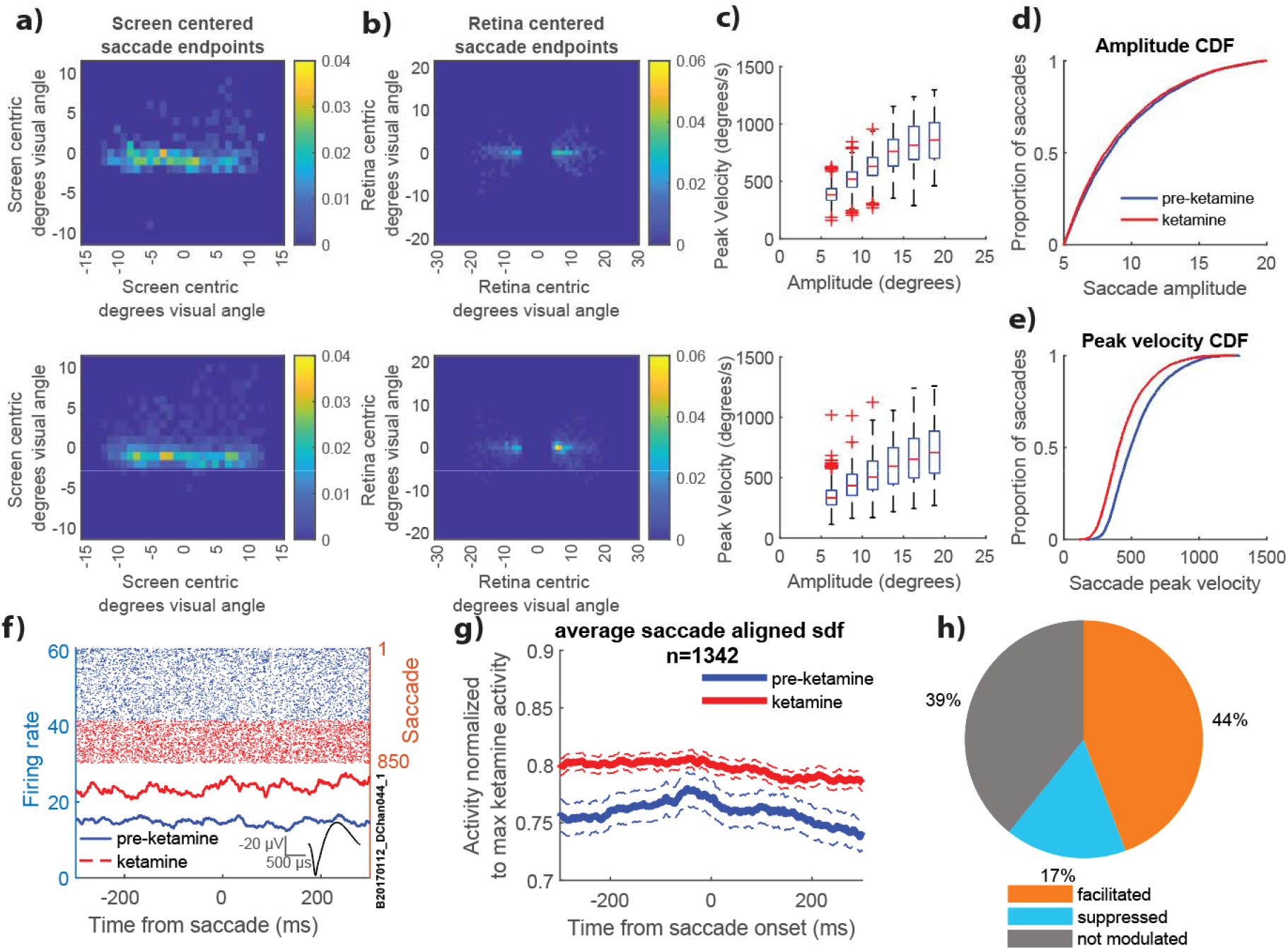
Saccadic activity is similar between pre and post injection, but with slower saccades; and neural activity is increased. a) Heatmap of end points for horizontal saccades between 5 and 20 degrees visual angle (DVA) from an example session in screen coordinates for pre-ketamine (above) and post-ketamine (below). b) Same as a) but in retina coordinates. c) The relationship between saccade amplitude and peak velocity follows a main sequence relationship for pre-ketamine (above) and post-ketamine (below). d) Cumulative density function for saccade amplitude starting at 5 DVA for pre-ketamine and ketamine trials. e) Cumulative density function for peak-velocity of saccades for pre-ketamine and ketamine trials. f) Example unit raster and spike density function (SDF) for saccade aligned activity during pre-ketamine (upper raster, blue, solid) and ketamine (lower raster, red, dashed) (inset: waveform) g) Average aligned activity for population, normalized to peak of ketamine SDF. Dashed lines represent 95% confidence intervals. h) Pie-chart depicting proportions of units that have firing rates that are significantly increased by ketamine, decreased by ketamine, or not affected.

### Effect of ketamine on single neuron responses

We recorded the responses of 1342 units (1134 monkey B, 208 monkey T) across 16 sessions (9 in monkey B and 7 in monkey T). We aligned unit activity to saccades and measured the firing rates during a pre-ketamine period from 300 to 200 ms before the saccade onset (**Fig 2f,g**). We used a rank sum analysis to determine if there was a change in firing rate. Monkey B had 698 units (61.6%) units modulated by ketamine, with 520 units that increased their firing rate, and 178 that decreased their firing rate, this is a ratio of ~3:1. Monkey T had 118 units (56.7%) units modulated by ketamine, with 74 units that increased their firing rate, and 44 that decreased their firing rate, this is a ratio of ~5:3. Overall 44% of units increased their firing rate, 17% decreased their firing rate, and 39% were not modulated (**Fig. 2h**). These results agree with previous reports on the effects of ketamine on neural responses in the LPFC (see (Roussy et al., 2021)).

To explore whether there was modulation of the saccadic signal, we first found units that were modulated by saccades using a rank sum test to compare firing rates for leftward and rightward saccades in 50 ms bins from −100 to +150 ms around saccade onset. Neurons with firing rates above baseline (150-100 ms before saccade onset) for a particular direction and a significant difference between directions were considered saccade modulated and analysed based on which direction they were significantly facilitated (**Fig. 3a-c**). In LPFC, we found 226 units (16%) were modulated by saccades (17% in monkey B and 15% in monkey T). Of these saccade-modulated units, 92 were selective for rightward, contraversive saccades (41%, 37% in monkey B and 66% in monkey T) and 134 units were selective for leftward, ipsiversive saccades (59%, 63% in monkey B and 34% in monkey T; **Fig. 3e**).

**Figure 3.**
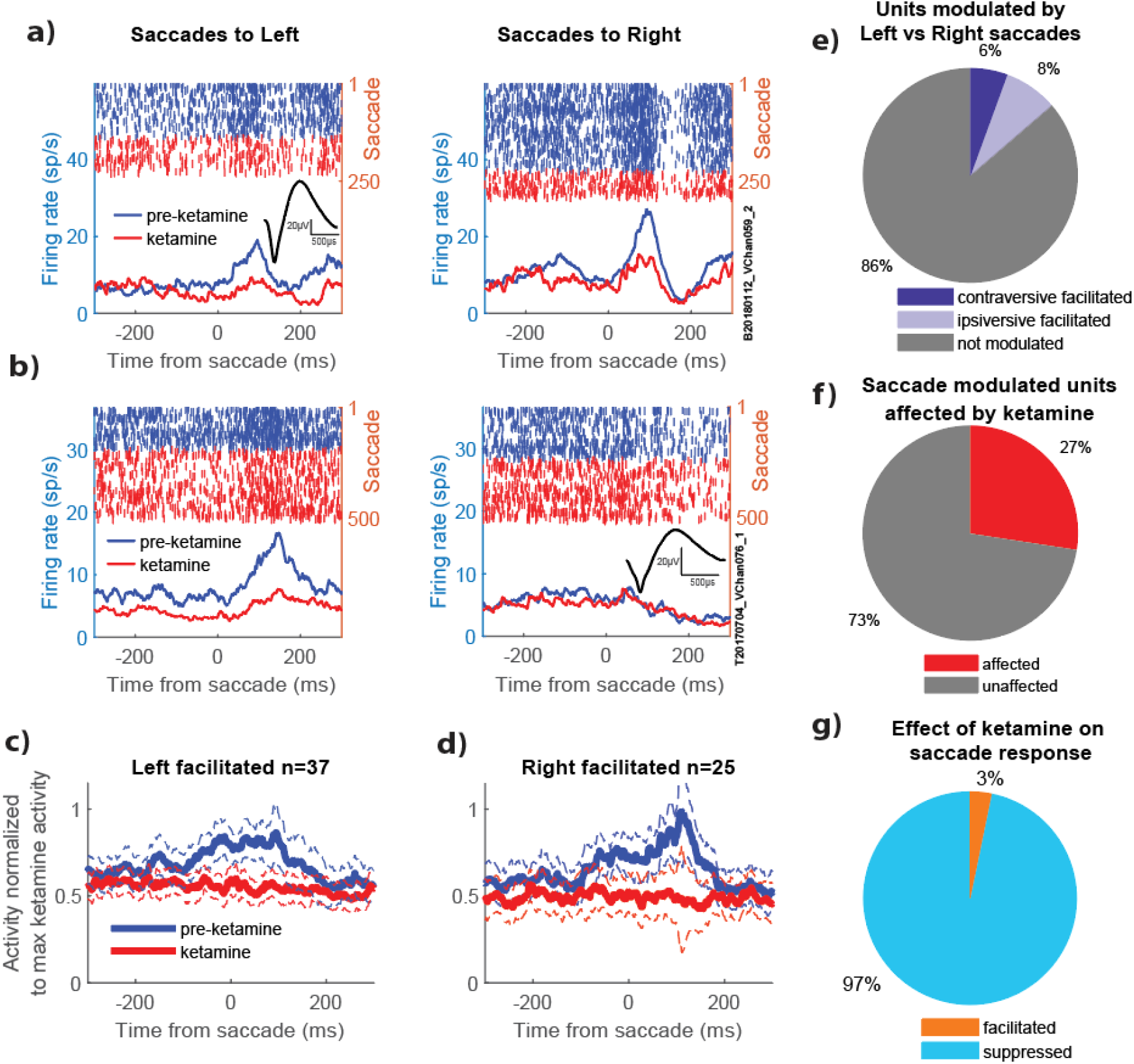
Ketamine reduces saccade related signals. a) Rasters and SDFs for an example unit from monkey B that increases firing rate after leftward saccades. This effect is decreased during saccades under ketamine. Average waveform inset. Scale bars represent 25 µV and 500 µs. b) Rasters and SDFs for an example unit from monkey T that increases firing rate after rightward saccades. This effect is decreased during saccades under ketamine. Average waveform inset. Scale bars represent 25 µV and 500 µs. c & d) Average population SDFs for left and right facilitated units during pre-ketamine and ketamine saccades. Dashed lines represent 95% confidence intervals. e) proportion of units with firing rates that are selectively facilitated by saccades to either the left or right. f) proportion of saccade modulated units that significantly increased or decreased by ketamine during the peak of the saccade modulation. g) most saccade modulated units that are affected by ketamine show a suppression of firing rate.

We then analysed these units for effects of ketamine during the 50ms bin of peak modulation. 62 units, or 27% of the saccade modulated units had significant effects of ketamine on saccade modulations (26% in monkey B and 38% in monkey T; **Fig. 3f**). In contrast to the trend of the majority of ketamine affected units having their firing rates enhanced (**Fig. 2g,h**), 97% of saccade selective ketamine modulated units showed response suppression, and only 3% show response enhancement (**Fig. 3g**). For the units that showed response suppression, only 36% also had firing rates reduced by ketamine at baseline (200 ms before the saccade), and 27% were actually facilitated at baseline under ketamine. This suggests that these effects are not simply a blanket reduction in firing rate caused by ketamine, but rather a reduction in the signal related to the saccade.

### Effect of ketamine on population codes

To analyse the effect of ketamine on saccade-related signals at the level of population codes, we pooled saccade-modulated units together to form a pseudopopulation. We then used a state-space analysis to interrogate how reliably the population of neurons represents the parameters of saccades. The latter is one of the conditions a CD must meet for the LPFC circuitry to predict the consequences of shifts in eye position and consequently in the retinal image after saccades, a process necessary for perceptual stability (Cavanaugh et al., 2016). For this analysis, we z-scored the firing rates and applied a Principal Components Analysis (PCA) during pre-ketamine, ketamine, and recovery periods, which started at 30 minutes post-injection to 1 hour post-injection.

We plotted trajectories in the state space using the first two PCs that explain 33.2% of the variance. In this low-dimensional space, the left and right saccade trajectories are well separated during the pre-ketamine period (**Fig. 4a,b**, blue lines). This is important because it indicates that the ensemble activity in the LPFC contains information about the direction of eye movement and can therefore predict changes in the retinal image after a saccade. Interestingly, after ketamine (red), the trajectories diverge from pre-ketamine (blue). This effect seems to partially recover 30-60 minutes after injection (purple). These details could also be appreciated in the PC1 and PC2 vs time trajectories (**Fig. 4c,d**).

**Figure 4.**
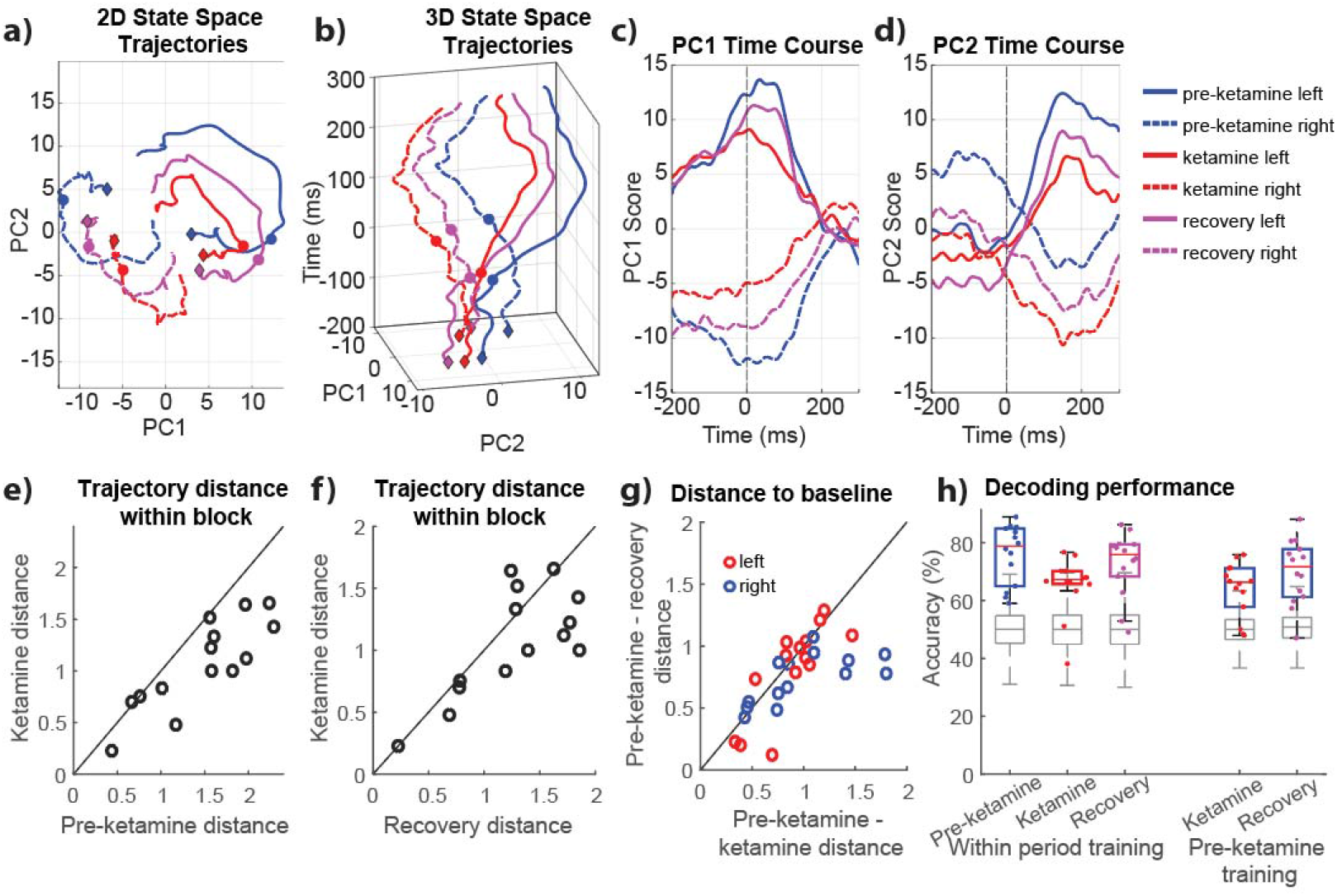
Saccade signals appear to start returning to pre-ketamine trajectories during recovery period. a) first two principal components (PCs) of pseudo-population of saccade modulated units across all sessions for left and right saccades (solid vs dashed). Diamonds indicate the first point, and circles indicate saccade onset. b) 3-dimensional trajectory of the first two PCs through time. c&d) First two PCs represented separately across time. Note that the recovery trajectories during the post saccade period are found between the pre-ketamine and ketamine trajectories. e) Scatterplot of the distance between the mean value of left and right saccade trajectories during the pre-ketamine and ketamine periods, where most points fall below the unity line, indicating that saccade trajectories are further apart before ketamine. f) Same as e, but comparing the ketamine to the recovery period, where points still trend to below the unity line. g) Comparison of distances to pre-ketamine trajectories for ketamine and recovery periods, where ketamine trajectories were significantly further away than recovery trajectories (see results). h) Saccade direction decoding accuracy when training within period, and the pre-ketamine decoder accuracy when tested on the ketamine and recovery periods.

If ketamine interferes with the CD saccade signal, it should impair the discriminability between the representations of left and right trajectories in the state space. We analysed the separability of representations corresponding to left and right saccades by measuring the Euclidian distance between the two direction trajectories for each period and session (n=14) for the 100ms period following saccade onset (**Fig. 4e**). Pre-ketamine trajectories had an average distance of 1.48 (SD = 0.59) and which was significantly higher than ketamine trajectories, which had a mean distance of 1.07 (SD = 0.44), using a t-test (*t*(26)=2.10, *p* = 0.045). Thus, ketamine decreased the discriminability by ~28%. The trajectories in the recovery period had a mean distance of 1.27 (SD =.50). This was not significantly different from either pre-ketamine or ketamine periods (*p* > 0.05 for both) (**Fig. 4f**). This recovery trajectories fall right in between, suggesting that although population codes were ‘recovering’ from the effects of ketamine, they did not do so to a full extent when we executed the last measurements of neural activity.

It is possible that during recovery, the representations landed in a different subspace from the pre-ketamine period, or that they were returning to the pre-ketamine subspace. To explore these alternatives, we calculated the Euclidean distance between pre-ketamine and ketamine trajectories (mean = 0.94 SD = 0.39) and pre-ketamine and recovery trajectories (mean = 0.77, SD = 0.30) for each direction (**Fig 4g**). There was an average shift towards pre-ketamine of 0.16 (SD = 0.31) across sessions during recovery, which was significantly different from zero using a paired t-test (*t*(15) = 2.60, *p* = 0.020). These results favor the hypothesis that, given enough recovery time, the representation would have landed in a subspace similar to pre-ketamine.

Finally, we hypothesized that ketamine would also reduce the ability of the LPFC population to discriminate left and right saccades in single trials. We used a support vector machine to decode saccade direction (left vs right) from ensembles of simultaneously recorded neurons, training and testing within each block for each session. During the pre-ketamine period, median accuracy was 78.8% (IQR = 19.9%) while median ketamine accuracy was 67.2% (IQR = 4.5%) and median recovery accuracy was in the middle at 75.9% (IQR = 11.1%, **Fig 4h**). All accuracies were significantly greater than the accuracies on shuffled data (median 50%) using a rank-sum test (*p*<< 0.05). We used a Kruskal-Wallis test to assess the effect of task period (pre-ketamine, ketamine, and recovery) on accuracy, which was significant (*H*(2, *n* = 28) =7.18, *p* = 0.028). Post-hoc rank-sum tests showed that ketamine accuracy was lower than both pre-ketamine (*z =* 2.18, *n* = 28, *p* = 0.029) and recovery (*z =* 2.27, *n* = 28, *p* = 0.023), and there was no significant difference between pre-ketamine and recovery (*z =* 0.90, *n* = 28, *p* = 0.370). These results demonstrate that ketamine impairs the decoding of saccade direction in single trials.

A second prediction derived from our previous analyses is that training a decoder in the pre-ketamine period and testing it in the ketamine period would produce lower performance than training the decoder in the pre-ketamine period and testing it in the recovery period. This is because population codes are more similar during pre-ketamine and recovery than during pre-ketamine and ketamine. Training during pre-ketamine and testing during ketamine produced a median performance of 66.3% (IQR = 13.3), while training pre-ketamine and testing during recovery produced a median performance of 71.7% (IQR = 16.4). The latter performance was significantly higher (signrank test, *z* = 2.12, *n* = 14, *p* =0.034).

## Discussion

Our study provides novel evidence that subanesthetic ketamine disrupts saccade-related corollary discharge (CD) signals in the primate lateral prefrontal cortex (LPFC). CD signals are critical for predictive coding: they allow the brain to predict the sensory consequences of eye movements and maintain perceptual stability across saccades. In both macaques we studied, ketamine selectively suppressed the activity of saccade-modulated LPFC neurons, impairing their ability to discriminate leftward from rightward saccades at the single-neuron and population levels. These effects were robust, reversible, and distinct from the general increase in baseline firing rates observed across most neurons. Thus, rather than simply elevating activity globally, ketamine specifically dampened CD signals, impairing the discriminability of self-generated actions.

### CD disruption as a mechanism of perceptual instability

CD signals provide the brain with predictive information about the sensory consequences of movements. In the oculomotor system, they allow cortical circuits to anticipate changes in the retinal image caused by saccades, ensuring perceptual stability despite rapid shifts in gaze. The presence of robust saccade-related signals in LPFC likely reflects this region’s critical role in processing saccadic information to update spatial representations and maintain cognitive maps of the environment (Bullock et al., 2017; Corrigan et al., 2023; Johnston et al., 2023). The LPFC receives extensive projections from both the Frontal Eye Fields (FEF) (Cohen et al., 2010; Monteon et al., 2013; Thompson et al., 1997), and the Superior Colliculus (SC), both involved in target selection and saccade generation, via the mediodorsal thalamus (Goldman□Rakic & Porrino, 1985). The LPFC also shows selectivity for visual features such as motion direction (Mendoza-Halliday et al., 2015; Zaksas & Pasternak, 2006), object categories (Freedman et al., 2001, 2002, 2003; Meyer et al., 2011), and abstract rules involving combinations of features and spatial locations (Rouzitalab et al., 2023; Wallis et al., 2001). The LPFC is positioned to integrate CD signals with ongoing cognitive processes, thereby achieving a stable representation of the world and making predictions about the future.

Our finding that ketamine selectively disrupts saccade-related signals while largely preserving baseline neural activity suggests a specific impact on CD processing. The observed suppression of direction-selective responses in 97% of saccade-modulated units indicates a breakdown in the precise encoding of eye movement parameters. While it is the case that approximately one-third of the units were also suppressed during their pre-ketamine period, most were not, and more than one-quarter were facilitated during the pre-ketamine period and still had suppressed saccade-related activity. This disruption could impair the LPFC’s ability to update spatial representations and maintain accurate internal models of the environment that remain robust to saccadic eye movements (Roussy et al., 2022).

### Implications for predictive coding

Predictive coding theories propose that cortical circuits continuously generate internal models of expected sensory input, which are compared against actual sensory signals. CD is a central component of this framework, as it conveys predictions derived from motor commands (Brower et al., 2019). One interesting finding in our study is that the population of eye movement selective neurons was able to discriminate between left and right saccades. Ketamine decreases such discriminability, which supports the argument that ketamine disrupts CD and predictions of their sensory consequences. During the recovery period, such changes gradually recovered to resemble the pre-ketamine period. Importantly, the decoding analyses showed that population activity in single trials was sufficient to discriminate the direction of the saccade, and this process was hindered by ketamine. These effects are unlikely to have motor origins since basic saccade metrics were relatively preserved. A similar result has been reported in rodent hippocampus where locomotion is relatively intact, but place cell responses are suppressed by ketamine administration (Masuda et al., 2023).

### Relevance to schizophrenia

Our findings provide strong support for theories linking NMDA receptor dysfunction, impaired CD, and psychotic symptoms (Thakkar & Rolfs, 2019; Yao et al., 2024). Schizophrenia patients often struggle to distinguish self-generated from externally generated experiences, and oculomotor tasks reveal specific CD deficits (Thakkar & Rolfs, 2019). Our results demonstrate how an NMDA antagonist can reproduce these deficits at the neuronal and circuit levels in primate LPFC. This convergence strengthens the translational validity of the ketamine model and suggests that CD disruption may be a fundamental mechanism contributing to hallucinations and cognitive fragmentation in schizophrenia (Yang et al., 2024).

Our results align with observations in schizophrenia patients, who show intact basic eye movements but impaired performance on tasks requiring integration of CD signals (Thakkar & Rolfs, 2019). The selective disruption of saccade-related neural responses we observed under ketamine mirrors the specific deficits in CD processing seen in schizophrenia, particularly in tasks requiring sequential eye movements or spatial updating (Richard et al., 2014; Thakkar et al., 2015). The disruption of CD signals in LPFC may contribute to the broader cognitive and perceptual disturbances observed in schizophrenia. When the brain cannot accurately track self-generated movements, it may lead to misattributions of agency and disrupted reality monitoring (Ford & Mathalon, 2012). Our findings suggest that NMDA receptor dysfunction in selective components of the LPFC circuit, as induced by ketamine, might be a key mechanism underlying these deficits. Future studies should investigate whether similar neural signatures are present in schizophrenia patients and whether selectively targeting NMDA receptor function might help restore normal corollary discharge processing.

The ketamine non-human primate model offers several advantages for investigating schizophrenia mechanisms (Gopinath et al., 2016; Ma et al., 2015; Maltbie et al., 2015; Roussy et al., 2021; Skoblenick et al., 2016; Wang et al., 2013). Unlike patient studies, it allows for direct single neuron recordings before and after symptom induction, enabling careful examination of cause-and-effect relationships in areas such as the LPFC, which are exclusively found in primates. The preservation of basic eye movements while disrupting higher-order processing closely mirrors the pattern seen in schizophrenia. In general, primate models could accelerate research by allowing rapid testing of therapeutic interventions targeting NMDA receptor function in brain areas that are homologous to human brain areas.

### Broader implications and future directions

By demonstrating that CD-related signals are selectively impaired by ketamine in the primate LPFC, our study highlights a neural mechanism that bridges motor control, sensory integration, and cognitive stability. Future work should examine how these disruptions interact with working memory representations and whether they generalize to other cortical regions and behavioral contexts. It will also be important to determine whether pharmacological or neuromodulatory interventions can restore CD fidelity, offering potential therapeutic avenues for disorders involving predictive coding deficits.

## Conclusion

In summary, we provide evidence that ketamine impairs CD signaling in primate LPFC, disrupting predictive coding mechanisms that stabilize perception across saccades. These findings identify a neural circuit vulnerability that may underlie ketamine’s dissociative effects and contribute to core symptoms of schizophrenia, while also advancing our understanding of how predictive signals support perception and cognition in the primate brain.

## Methods

Two adult male rhesus macaques (Macaca mulatta) were used in this experiment (age: 10, 9 years; weight: 12, 10 kg). Animal care, handling and procedures were pre-approved by the University of Western Ontario Animal Care Committee. This approval ensure that standards for the ethical use of animals were followed as set by provincial (Ontario Animals in Research Act), federal (Canadian Council on Animal Care) and other regulatory bodies e.g., CIHR/NSERC). Regular assessments for physical and physiological well-being of the animals were carried out by veterinarians, registered veterinary technicians and researchers.

### Experimental set-up

The set-up and procedures are detailed in Roussy et al. (Roussy et al., 2021), and described briefly here. The monkey was seated in a primate chair in front of a 27” LCD monitor positioned 80cm away with a refresh rate of 75 Hz. Eye-position was recorded at 500 Hz (EyeLink 1000, SR Research). A custom computer program controlled the stimulus presentation, reward schedule and synchronized eye behavior and task events. Before recordings were carried out, a custom fit PEEK cranial implant with connections for headpost and recording equipment was implanted (Neuronitek, see Blonde et al., (Blonde et al., 2018)). The head post was attached to the chair to immobilize the head, and a two-axis joystick was used for virtual navigation.

### Task

The task is covered in detail in Roussy et al. (2021), but briefly, the task took place in a virtual environment where target locations in a circular arena were arranged in a 3 × 3 grid and spaced ~.5 s apart, as movement speed was fixed. Task periods were separated into pre-ketamine baseline period, post-injection ketamine period (0-30 minutes) and recovery period (30-60 minutes post-injection).

### Microelectrode array

All surgical procedures were carried out under general anesthesia induced by ketamine and maintained using isoflurane and propofol. Two Utah arrays with 96 channels in a 10×10 grid with.4 mm spacing and 1.5 mm length (Blackrock Microsystems) were chronically implanted on either side of the principal sulcus, just anterior to the knee of the arcuate sulcus, in the lateral prefrontal cortex (LPFC). Targeting of the craniotomy was achieved using brain navigation software Brainsight (Rogue Research Inc.). Arrays were placed and impacted ~1.5mm into the cortex.

### Neuronal recordings

Neurophysiological data was recorded using a Cerebus Neural Signal Processor (Blackrock Microsystems) via a Cereport adapter at 30 kHz. Spikes were detected online via a threshold crossing set at 3.4 standard deviations of the signal for each channel. Spikes were semi-automatically sorted using Plexon Offline Sorter (Plexon Inc.). We collected behavioural data across 16 ketamine-WM sessions (9 in monkey B, 7 in monkey T), yielding 1652 single units. Only units with a firing rate of at least.5 spikes/s across the session were used. If a unit’s average firing rate during the last 100 trials of the recovery period decreased to less than 1/3 the average rate over the first up to 100 trials of the pre-ketamine period, or increased to more than 3x that rate, the unit was excluded based on an assumption of poor isolation during one of the periods. This left 1342 units (1134 in monkey B and 208 in monkey T).

### Ketamine Injection

Animals were trained to receive injections while seating in the primate chair within the experimental set-up. Ketamine was injected intramuscularly (0.25, 0.4, 0.8 mg/kg) into the hamstring muscles by a registered veterinary technician. The ketamine doses were titrated so they did not induce visible behavioural changes in the animals such as nystagmus or somnolence. Ketamine injections were spaced at least 2 days apart to allow for washout of the drug (Zanos et al., 2018).

### Behaviour performance

Monkeys had to navigate to the cued location, and the trial would end once they reached it, or after 10 s, which was considered a miss. We calculated performance based on how many trials ended because the monkey made it to the correct location.

### Eye-movement behaviour

We recorded eye-position and calculated the onset and offsets of saccade using the method described in Corrigan et al. (Corrigan et al., 2017). Briefly, we used an adaptive acceleration threshold to identify time periods with saccades, found the peak velocity, and then calculated the deviation in direction from the peak velocity direction for each point going forward and backward in time to find when the movement transitioned to a foveation. The first point to point change with a speed less than the greater of 30°/s or 1/5 of the peak velocity where there was a direction deviation from peak velocity direction of more than 60°, or the third point with a deviation >20°, or just a point to point change of >60° or the third point to point change of >20° was considered the onset or offset of the saccade.

### Neural analyses

To assess stationarity of neurons, we set a threshold where units had to have changed firing rates by a maximum of a factor of three (x3 or 1/3) between the pre-ketamine period and the recovery period. To measure the effects ketamine, we aligned spikes to saccades but then analysed the period 300 to 200 ms before the saccade by calculating the average firing rate pre- and post-injection. To determine whether a saccade was modulated differentially by leftward or rightward saccades we first matched saccades based on both landing location and amplitude. We binned visual space into bins of 4 degrees visual angle, and saccade amplitude into bins of 7 degrees visual angle starting at 5 degrees. This way similar amplitude saccades would land on similar parts of the screen and the objects foveated would not drive differences between saccade directions as might happen with a view cell (Corrigan et al., 2023). We ran this matching for directions and for task periods, so that there were the same number of right and left saccades of similar amplitudes and positions for each task period. To determine if a neuron was modulated by direction, we used two-tailed ranksum tests to compare firing rates in 50ms bins from −100:150 ms around the saccade onset and used an alpha of.05. To classify this modulation as facilitation, we again used a ranksum but then compared the population with the higher firing rates against the baseline firing rate of the combined populations from −150:−100 ms before the saccade. Here we used a one-sided alpha of.02, and the peri-saccadic bin with the greatest difference from the baseline was considered the peak modulation bin. So, to be modulated, the neuron needed to exhibit an elevated firing rate above baseline in a particular bin and above the firing rate for the opposing direction for that bin.

Once we found saccade modulated neurons that had a peri-saccadic response that was differentially affected by direction, we then compared the firing rates for the saccades the neuron was selective for during the peak modulation bin to the same bin for the same direction saccades post-ketamine injection using a two-sided ranksum test to identify neurons that had saccade modulated signals affected by ketamine.

For state-space analyses, we convolved the firing rate of each trial with an alpha function with a rise of 1ms and a decay of 20 ms to approximate the shape of a postsynaptic potential (Hanes & Schall, 1996). These spike density functions were then averaged for each direction and period and smoothed with a 20 ms moving average filter and finally z-scored across all directions and periods from −200:300 ms relative to saccade. PCA was run on these 6 populations to create the visualizations, but analyses were calculated based on the Euclidian distances between the average position for the first 100 ms following saccade onset.

For decoding analyses, we used a support vector machine (SVM;LIBSVM (Chang & Lin, 2011)) with a linear kernel. For each session that had at least 3 saccade modulated neurons, calculated the firing rates for the period from −50 to 150 ms after saccade onset, then normalized them between 0 and 1 and ran a 5 k-fold train test split for right and left saccades in each period. To compute the chance decoding, we shuffled the labels and re-ran the analysis 100 times. We also trained the SVM on the whole pre-ketamine period, and then tested it on the ketamine and recovery periods, as well as shuffled labels to get chance decoding.

## Acknowledgements

We thank registered veterinary technicians Kim Thomaes and Rhonda Kersten from the University of Western Ontario for their assistance in surgery and animal care; Guillaume Doucet from the University of Ottawa for technical assistance related to Unreal Development Kit; Kevin Barker from Neuronitek for engineering equipment for our experiments; Jonathan C. Lau from the Division of Neurosurgery, University Hospital for providing advice regarding surgery and surgical planning. This work was supported by Natural Sciences and Engineering Research Council of Canada (NSERC) Doctoral Scholarship (to B.W.C. & M.R.) and Canadian Institute of Health Research (CIHR) Postdoctoral Fellowship (to B.W.C); Ontario Graduate Scholarship; Jonathan & Joshua Memorial Graduate Scholarship in Mental Health Research. Chrysalis Foundation (London, Ontario). LP acknowledges salary support from the Tanna Schulich Endowment Chair for Neuroscience and Mental Health. CIHR Project Grant; NSERC; BrainsCAN, NEURONEX Brain Initiative (ref. FL6GV84CKN57) grants (to J.C.M.).

## Author Contributions

BWC analyzed data, interpreted data and wrote the manuscript. MR planned the study, designed the task, trained animals, planned and conducted surgeries, collected data, conducted data preprocessing and contributed to manuscript revisions. RL trained animals, planned and conducted surgeries, collected data, conducted data preprocessing and contributed to manuscript revisions. RAG planned and conducted surgeries, developed code for data preprocessing and contributed to experimental design. JDS contributed to manuscript revisions. AJS planned and conducted surgeries. LP planned the study and contributed to manuscript revisions. JCM planned the study, designed the task, planned and conducted surgeries, interpreted the data and wrote the manuscript.

## Data availability

Data supporting the findings of this study is available from the corresponding authors on reasonable request and will be fulfilled by BWC

## Code availability

Custom code for data analysis was written in MATLAB and is available from the corresponding authors on reasonable request and will be fulfilled by BWC.

## Citations

Abram, S. V., Roach, B. J., Fryer, S. L., Calhoun, V. D., Preda, A., van Erp, T. G. M., Bustillo, J. R., Lim, K. O., Loewy, R. L., Stuart, B. K., Krystal, J. H., Ford, J. M., & Mathalon, D. H. (2022). Validation of ketamine as a pharmacological model of thalamic dysconnectivity across the illness course of schizophrenia. Molecular Psychiatry, 27(5), 2448–2456. 10.1038/s41380-022-01502-0

Anticevic, A., Haut, K., Murray, J. D., Repovs, G., Yang, G. J., Diehl, C., McEwen, S. C., Bearden, C. E., Addington, J., Goodyear, B., Cadenhead, K. S., Mirzakhanian, H., Cornblatt, B. A., Olvet, D., Mathalon, D. H., McGlashan, T. H., Perkins, D. O., Belger, A., Seidman, L. J., … Cannon, T. D. (2015). Association of Thalamic Dysconnectivity and Conversion to Psychosis in Youth and Young Adults at Elevated Clinical Risk. JAMA Psychiatry, 72(9), 882–891. 10.1001/JAMAPSYCHIATRY.2015.0566

Blonde, J. D., Roussy, M., Luna, R., Mahmoudian, B., Gulli, R. A., Barker, K. C., Lau, J. C., & Martinez-Trujillo, J. C. (2018). Customizable cap implants for neurophysiological experimentation. Journal of Neuroscience Methods, 304, 103–117. 10.1016/j.jneumeth.2018.04.016

Brower, R., Wang, H. R., Bansal, S., & Joiner, W. M. (2019). Using Corollary Discharge and Predictive Coding to Understand False Sensations and Beliefs. Biological Psychiatry: Cognitive Neuroscience and Neuroimaging, 4(9), 770–772. 10.1016/j.bpsc.2019.07.004

Bullock, K. R., Pieper, F., Sachs, A. J., & Martinez-Trujillo, J. C. (2017). Visual and presaccadic activity in area 8Ar of the macaque monkey lateral prefrontal cortex. Journal of Neurophysiology, 118(1), 15–28. 10.1152/jn.00278.2016

Busch, A., Roussy, M., Luna, R., Leavitt, M. L., Mofrad, M. H., Gulli, R. A., Corrigan, B., Mináč, J., Sachs, A. J., Palaniyappan, L., Muller, L., & Martinez-Trujillo, J. C. (2024). Neuronal activation sequences in lateral prefrontal cortex encode visuospatial working memory during virtual navigation. Nature Communications, 15(1), 4471. 10.1038/s41467-024-48664-9

Busettini, C., & Frölich, M. A. (2014). Effects of mild to moderate sedation on saccadic eye movements. Behavioural Brain Research, 272, 286–302. 10.1016/j.bbr.2014.07.012

Cavanaugh, J., Berman, R. A., Joiner, W. M., & Wurtz, R. H. (2016). Saccadic Corollary Discharge Underlies Stable Visual Perception. Journal of Neuroscience, 36(1), 31–42. 10.1523/JNEUROSCI.2054-15.2016

Chang, C.-C., & Lin, C.-J. (2011). LIBSVM: A Library for Support Vector Machines. ACM Transactions on Intelligent Systems and Technology, 2(3), 1–27. 10.1145/1961189.1961199

Cohen, J. Y., Crowder, E. A., Heitz, R. P., Subraveti, C. R., Thompson, K. G., Woodman, G. F., & Schall, J. D. (2010). Cooperation and Competition among Frontal Eye Field Neurons during Visual Target Selection. Journal of Neuroscience, 30(9), 3227–3238. 10.1523/JNEUROSCI.4600-09.2010

Corrigan, B. W., Gulli, R. A., Doucet, G., Mahmoudian, B., Abbass, M., Roussy, M., Luna, R., Sachs, A. J., & Martinez-Trujillo, J. C. (2023). View cells in the hippocampus and prefrontal cortex of macaques during virtual navigation. Hippocampus, 33(5), 573–585. 10.1002/HIPO.23534

Corrigan, B. W., Gulli, R. A., Doucet, G., & Martinez-Trujillo, J. C. (2017). Characterizing eye movement behaviors and kinematics of non-human primates during virtual navigation tasks. Journal of Vision, 17(12), 15. 10.1167/17.12.15

Crawford, J. D., Martinez-Trujillo, J. C., & Klier, E. M. (2003). Neural control of three-dimensional eye and head movements. Current Opinion in Neurobiology, 13(6), 655–662. 10.1016/J.CONB.2003.10.009

Ford, J. M., & Mathalon, D. H. (2012). Anticipating the future: automatic prediction failures in schizophrenia. International Journal of Psychophysiology□: Official Journal of the International Organization of Psychophysiology, 83(2), 232–239. 10.1016/j.ijpsycho.2011.09.004

Freedman, D. J., Riesenhuber, M., Poggio, T., & Miller, E. K. (2001). Categorical Representation of Visual Stimuli in the Primate Prefrontal Cortex. Science, 291(5502), 312– 316. 10.1126/SCIENCE.291.5502.312

Freedman, D. J., Riesenhuber, M., Poggio, T., & Miller, E. K. (2002). Visual Categorization and the Primate Prefrontal Cortex: Neurophysiology and Behavior. Journal of Neurophysiology, 88(2), 929–941. 10.1152/jn.2002.88.2.929

Freedman, D. J., Riesenhuber, M., Poggio, T., & Miller, E. K. (2003). A Comparison of Primate Prefrontal and Inferior Temporal Cortices during Visual Categorization. Journal of Neuroscience, 23(12), 5235–5246. 10.1523/JNEUROSCI.23-12-05235.2003

Funahashi, S., Bruce, C. J., & Goldman-Rakic, P. S. (1991). Neuronal activity related to saccadic eye movements in the monkey’s dorsolateral prefrontal cortex. Journal of Neurophysiology, 65(6), 1464–1483. 10.1152/jn.1991.65.6.1464

Goldman-Rakic, P.. (1995). Cellular basis of working memory. Neuron, 14(3), 477–485. 10.1016/0896-6273(95)90304-6

Goldman□Rakic, P. S., & Porrino, L. J. (1985). The primate mediodorsal (MD) nucleus and its projection to the frontal lobe. The Journal of Comparative Neurology, 242(4), 535–560. 10.1002/CNE.902420406

Gopinath, K., Maltbie, E., Urushino, N., Kempf, D., & Howell, L. (2016). Ketamine-induced changes in connectivity of functional brain networks in awake female nonhuman primates: a translational functional imaging model. Psychopharmacology, 233(21–22), 3673–3684. 10.1007/S00213-016-4401-Z

Hanes, D. P., & Schall, J. D. (1996). Neural Control of Voluntary Movement Initiation. Science, 274(5286), 427–430. 10.1126/science.274.5286.427

Johnston, R., Abbass, M., Corrigan, B., Gulli, R., Martinez-Trujillo, J., & Sachs, A. (2023). Decoding spatial locations from primate lateral prefrontal cortex neural activity during virtual navigation. Journal of Neural Engineering, 20(1). 10.1088/1741-2552/acb5c2

Krystal, J. H., D’Souza, D. C., Mathalon, D., Perry, E., Belger, A., & Hoffman, R. (2003). NMDA receptor antagonist effects, cortical glutamatergic function, and schizophrenia: toward a paradigm shift in medication development. Psychopharmacology 2003 169:3, 169(3), 215–233. 10.1007/S00213-003-1582-Z

Krystal, J. H., Karper, L. P., Seibyl, J. P., Freeman, G. K., Delaney, R., Bremner, J. D., Heninger, G. R., Bowers, M. B., & Charney, D. S. (1994). Subanesthetic effects of the noncompetitive NMDA antagonist, ketamine, in humans. Psychotomimetic, perceptual, cognitive, and neuroendocrine responses. Archives of General Psychiatry, 51(3), 199–214. 10.1001/ARCHPSYC.1994.03950030035004

Ma, L., Skoblenick, K., Seamans, J. K., & Everling, S. (2015). Ketamine-Induced Changes in the Signal and Noise of Rule Representation in Working Memory by Lateral Prefrontal Neurons. The Journal of Neuroscience□: The Official Journal of the Society for Neuroscience, 35(33), 11612–11622. 10.1523/JNEUROSCI.1839-15.2015

Maltbie, E., Gopinath, K., Urushino, N., Kempf, D., & Howell, L. (2015). Ketamine-induced brain activation in awake female nonhuman primates: a translational functional imaging model. Psychopharmacology, 233(6), 961. 10.1007/S00213-015-4175-8

Masuda, F. K., Aery Jones, E. A., Sun, Y., & Giocomo, L. M. (2023). Ketamine evoked disruption of entorhinal and hippocampal spatial maps. Nature Communications, 14(1), 6285. 10.1038/S41467-023-41750-4

Mendoza-Halliday, D., Torres, S., & Martinez-Trujillo, J. (2015). Working Memory Representations of Visual Motion along the Primate Dorsal Visual Pathway. In Mechanisms of Sensory Working Memory (pp. 159–169). Elsevier. 10.1016/B978-0-12-801371-7.00013-2

Meyer, T., Qi, X. L., Stanford, T. R., & Constantinidis, C. (2011). Stimulus Selectivity in Dorsal and Ventral Prefrontal Cortex after Training in Working Memory Tasks. Journal of Neuroscience, 31(17), 6266–6276. 10.1523/JNEUROSCI.6798-10.2011

Moghaddam, B., & Javitt, D. (2011). From Revolution to Evolution: The Glutamate Hypothesis of Schizophrenia and its Implication for Treatment. Neuropsychopharmacology, 37(1), 4. 10.1038/NPP.2011.181

Monteon, J. A., Wang, H., Martinez-Trujillo, J., & Crawford, J. D. (2013). Frames of reference for eye–head gaze shifts evoked during frontal eye field stimulation. European Journal of Neuroscience, 37(11), 1754–1765. 10.1111/EJN.12175

Pouget, P., Daye, P., & Paré, M. (2025). Cognitive and kinematic markers of ketamine effects in behaving non-human primates. European Journal of Pharmacology, 987. 10.1016/j.ejphar.2024.177185

Richard, A., Churan, J., Whitford, V., O’Driscoll, G. A., Titone, D., & Pack, C. C. (2014). Perisaccadic Perception of Visual Space in People with Schizophrenia. Journal of Neuroscience, 34(14), 4760–4765. 10.1523/JNEUROSCI.4744-13.2014

Rösler, L., Rolfs, M., van der Stigchel, S., Neggers, S. F. W., Cahn, W., Kahn, R. S., & Thakkar, K. N. (2015). Failure to use corollary discharge to remap visual target locations is associated with psychotic symptom severity in schizophrenia. Journal of Neurophysiology, 114(2), 1129–1136. 10.1152/jn.00155.2015

Roussy, M., Corrigan, B., Luna, R., Gulli, R. A., Sachs, A. J., Palaniyappan, L., & Martinez-Trujillo, J. C. (2022). Stable working memory and perceptual representations in macaque lateral prefrontal cortex during naturalistic vision. The Journal of Neuroscience, 42(44), JN-RM-0597-22. 10.1523/JNEUROSCI.0597-22.2022

Roussy, M., Luna, R., Duong, L., Corrigan, B., Gulli, R. A., Nogueira, R., Moreno-Bote, R., Sachs, A. J., Palaniyappan, L., & Martinez-Trujillo, J. C. (2021). Ketamine disrupts naturalistic coding of working memory in primate lateral prefrontal cortex networks. Molecular Psychiatry, 26(11), 6688–6703. 10.1038/s41380-021-01082-5

Rouzitalab, A., Boulay, C. B., Park, J., Martinez-Trujillo, J. C., & Sachs, A. J. (2023). Ensembles code for associative learning in the primate lateral prefrontal cortex. Cell Reports, 42(5), 112449. 10.1016/j.celrep.2023.112449

Selemon, L. D., & Goldman-Rakic, P. S. (1988). Common cortical and subcortical targets of the dorsolateral prefrontal and posterior parietal cortices in the rhesus monkey: evidence for a distributed neural network subserving spatially guided behavior. Journal of Neuroscience, 8(11), 4049–4068. 10.1523/JNEUROSCI.08-11-04049.1988

Skoblenick, K. J., Womelsdorf, T., & Everling, S. (2016). Ketamine Alters Outcome-Related Local Field Potentials in Monkey Prefrontal Cortex. Cerebral Cortex, 26(6), 2743–2752. 10.1093/CERCOR/BHV128

Subramanian, D., Alers, A., & Sommer, M. A. (2019). Corollary Discharge for Action and Cognition. In Biological Psychiatry: Cognitive Neuroscience and Neuroimaging (Vol. 4, Issue 9, pp. 782–790). 10.1016/j.bpsc.2019.05.010

Thakkar, K. N., & Rolfs, M. (2019). Disrupted Corollary Discharge in Schizophrenia: Evidence From the Oculomotor System. In Biological Psychiatry: Cognitive Neuroscience and Neuroimaging (Vol. 4, Issue 9, pp. 773–781). Elsevier. 10.1016/j.bpsc.2019.03.009

Thakkar, K. N., Schall, J. D., Heckers, S., & Park, S. (2015). Disrupted saccadic corollary discharge in schizophrenia. Journal of Neuroscience, 35(27), 9935–9945. 10.1523/JNEUROSCI.0473-15.2015

Thompson, K. G., Bichot, N. P., & Schall, J. D. (1997). Dissociation of Visual Discrimination From Saccade Programming in Macaque Frontal Eye Field. Journal of Neurophysiology, 77(2), 1046–1050. 10.1152/jn.1997.77.2.1046

Wallis, J. D., Anderson, K. C., & Miller, E. K. (2001). Single neurons in prefrontal cortex encode abstract rules. Nature, 411(6840), 953–956. 10.1038/35082081

Wang, M., Yang, Y., Wang, C.-J., Gamo, N. J., Jin, L. E., Mazer, J. A., Morrison, J. H., Wang, X.-J., & Arnsten, A. F. T. (2013). NMDA Receptors Subserve Persistent Neuronal Firing during Working Memory in Dorsolateral Prefrontal Cortex. Neuron, 77(4), 736–749. 10.1016/J.NEURON.2012.12.032

Weinberger, D. R., Berman, K. F., & Zec, R. F. (1986). Physiologic dysfunction of dorsolateral prefrontal cortex in schizophrenia. I. Regional cerebral blood flow evidence. Archives of General Psychiatry, 43(2), 114–124. 10.1001/ARCHPSYC.1986.01800020020004

Yang, F., Zhu, H., Cao, X., Li, H., Fang, X., Yu, L., Li, S., Wu, Z., Li, C., Zhang, C., & Tian, X. (2024). Impaired motor-to-sensory transformation mediates auditory hallucinations. PLOS Biology, 22(10), e3002836. 10.1371/JOURNAL.PBIO.3002836

Yao, B., Rolfs, M., Slate, R., Roberts, D., Fattal, J., Achtyes, E. D., Tso, I. F., Diwadkar, V. A., Kashy, D., Bao, J., & Thakkar, K. N. (2024). Abnormal Oculomotor Corollary Discharge Signaling as a Trans-diagnostic Mechanism of Psychosis. Schizophrenia Bulletin, 50(3), 631–641. 10.1093/SCHBUL/SBAD180

Zaksas, D., & Pasternak, T. (2006). Directional Signals in the Prefrontal Cortex and in Area MT during a Working Memory for Visual Motion Task. Journal of Neuroscience, 26(45), 11726–11742. 10.1523/JNEUROSCI.3420-06.2006

Zanos, P., Moaddel, R., Morris, P. J., Riggs, L. M., Highland, J. N., Georgiou, P., Pereira, E. F. R., Albuquerque, E. X., Thomas, C. J., Zarate, C. A., & Gould, T. D. (2018). Ketamine and Ketamine Metabolite Pharmacology: Insights into Therapeutic Mechanisms. Pharmacological Reviews, 70(3), 621. 10.1124/PR.117.015198

